# Identification and Validation of N-Acetylputrescine in Combination With Non-Canonical Clinical Features As a Parkinson’s Disease Biomarker Panel

**DOI:** 10.1101/2021.07.23.453542

**Authors:** Kuan-Wei Peng, Allison Klotz, Arcan Guven, Unnati Kapadnis, Shobha Ravipaty, Vladimir Tolstikov, Vijetha Vemulapalli, Leonardo O. Rodrigues, Hongyan Li, Mark D. Kellogg, Farah Kausar, Linda Rees, Rangaprasad Sarangarajan, Birgitt Schüle, William Langston, Paula P. Narain, Niven R. Narain, Michael A. Kiebish

## Abstract

Parkinson’s disease is a progressive neurodegenerative disorder in which loss of dopaminergic neurons in the substantia nigra results in a clinically heterogeneous group with variable motor and non-motor symptoms with a degree of misdiagnosis. Only 3-5% of sporadic Parkinson’s patients present with genetic abnormalities, thus environmental, metabolic, and other unknown causes contribute to the pathogenesis of Parkinson’s disease, which highlights the critical need for biomarkers. There could be a significant clinical benefit to treating Parkinson’s disease at the earliest stage and identify at-risk populations once disease-modifying treatments are available. In the present study, we prospectively collected and analyzed plasma samples from 201 Parkinson’s disease patients and 199 age-matched non-diseased controls. Multiomic and Bayesian artificial intelligence analysis of molecular and clinical data identified the diagnostic utility of N-acetyl putrescine (NAP) in combination with smell (B-SIT), depression/anxiety (HADS), and acting out dreams (RBD1Q) clinical measurements. The clinical and biomarker panel demonstrated an area under the curve, AUC = 0.9, positive predictive value, PPV = 0.91, and negative predictive value, NPV = 0.66 utilizing all four variables. The assessed diagnostic panel demonstrates combinatorial utility in diagnosing Parkinson’s disease, allowing for an integrated interpretation of disease pathophysiology and highlighting the use of multi-tiered panels in neurological disease diagnosis.

## 1. Introduction

Parkinson’s Disease (PD) is a progressive neurological disorder characterized by motor features including tremors, bradykinesia, muscle rigidity, and postural instability, as well as non-motor features such as loss of sense of smell, sleep/REM disorder, and autonomic dysfunctions which can include constipation, urinary problems, changes in heart rate variability, psychiatric disturbances with anxiety, and depression as well as cognitive decline [1]. Pathologically, PD is defined by dopaminergic neuronal loss in the substantia nigra pars compacta (SN), and intracellular inclusions called Lewy bodies (LB) in the neurons of affected brain regions **[2, 3]**. Abnormal handling of misfolded proteins by the ubiquitin–proteasome and the autophagy–lysosomal systems **[4, 5]**, increased oxidative stress, mitochondrial dysfunction, and inflammation, well-described mechanisms involved in the pathogenesis of PD [5, 6].

The misdiagnosis rate of patients with PD in the clinical setting can be as high as 25%-42% [7] in the early stage of the disease. **[8–10]**. A molecular diagnostic test, which can be used to identify those with early stages of PD is a critical unmet need. Reliable diagnostic biomarkers are essential for identifying populations at risk and those that are pathologically susceptible to disease impetus. This provides an opportunity for an early and accurate diagnosis to predict the disease occurrence and progression. The diagnostic biomarker can be used as an objective tool to characterize evaluation indicators stratifying normal and pathogenic biological processes **[11]**.

Significant progress has been made in uncovering the complex molecular mechanisms exploited in the pathogenesis of PD. The emergence of several omics techniques, such as transcriptomics, proteomics, and metabolomics, have played a key role in identifying novel pathways associated with dopaminergic neurodegeneration, global system physiological changes, and subsequently PD, which include mitochondrial and proteasomal function as well as synaptic neurotransmission **[10]**. Additionally, these unbiased techniques, particularly in the brain regions that are uniquely associated with the disease, have greatly enhanced our ability to identify novel pathways, such as axon-guidance, potentially involved in PD pathogenesis **[12, 13].** To date, there has been extensive focus on the genetic etiology of disease [3]. **In contrast**, multi-omic analysis provides broader connectivity to adaptive and environmental sequalae that drive disease phenotypic effectors. Metabolomics assessment, which comprises the broadest capture of integrated biochemical assessment using mass spectrometry or NMR technology, provides a comprehensive view of metabolites tied to the biological phenotype.

Interestingly in PD patients, putrescine levels are increased in cerebrospinal fluid (CSF), whereas the concentration of spermidine is reduced compared to controls **[15]**. Putrescine is a polyamine that belongs to the category of ubiquitous small polycations that ionically bind to various negatively charged molecules and have many functions, mostly linked to cell growth, survival, and proliferation **[14]**. Examples of polyamines are putrescine, spermidine and spermine, whose levels are stringently regulated in the human body. Based on partial polyamine data previously reported **[15]** and our metabolomics platform outcome, we investigated a prospective PD cohort clinically and evaluated polyamine metabolic changes analytically. Further, we integrated clinical features identified by our Bayesian analysis to be causally associated with clinical outcomes. The use of these clinical features as a phenotypic readout complimented the molecular biomarker analysis. The combination of the CLIA validated assay of N-Acetylputrescine (NAP) and non-canonical clinical features of Hospital Anxiety and Depression Scale (HADS) and REM Sleep Behavior Disorder Single-Question Screen score (RBD1Q) demonstrated diagnostic utility [16–18]. This panel might provide broad utility for PD diagnosis and represents the integration of both clinical and molecular presentation of the disease.

## 2. Materials and Methods

### 2.1 Materials

N-Acetylputrescine reference standard, Bovine Serum Albumin (BSA), Trichloroacetic acid (TCA), Formic Acid (FA) and Isobutyl Chloroformate (IBCF) were obtained from Sigma (St. Louis, MO, USA). Optima LC/MS Grade of Acetonitrile, Water, and Sodium carbonate were purchased from Fisher Scientific (Pittsburgh, PA,USA). Human K_2_-EDTA Plasma was supplied by BIOIVT (Westbury, NY, USA)

### 2.2 Plasma Sample Collection

K_2_-EDTA plasma samples and clinical data from PD patients and controls were obtained from Parkinson’s Institute and Clinical Center (Sunnyvale, CA., USA). All study subject information shall remain de-identified. Samples were collected within the following range of self-reported fasting time: not less than 4 hours, not more than 8 hours. Non-diseased controls (n=199) and Parkinson’s Disease patients (n=201) were analyzed, with Hoehn and Yahr (H&Y) average scale of 2.1 and Unified Parkinson’s Disease rating scale (UPDRS) score from 10 to 115 (Table 1). The non-diseased control group included 125 males and 72 females. The PD patient cohort included 103 males and 91 females. All volunteers participating in this study gave their informed consent for inclusion before they participated in the study. Research use of the samples was conducted by the terms outlined within the informed consent form and the terms set forth therein, along with the tenets of the Declaration of Helsinki and its later amendments or comparable ethical standards.

**Table 1.** Patient Demographics.

|  | **Non-Disease** | **Parkinson's Disease** |
| --- | --- | --- |
| **Total Patients** | 197 | 194 |
| **Male** | 125 | 103 |
| **Female** | 72 | 91 |
| **Age, mean years (SD)** | 65.4 (9.6) | 64.7 (9.1) |
| **H&Y Stage** |  |  |
| **Mean ± SD** | N/A | 2.1 ± 0.7 |
| **Case (n)** | N/A | I(43), II(127),III (22), IV (7), V(1) |
| **UPDRS mean, (SD), (range)** | N/A | 40.4, (16.9), (10-115) |
| **UPDRS-III mean, (SD), (range)** | N/A | 25.6,(11.0), (1-69) |

### 2.3 Sample Preparation of NAP in Plasma Extraction and Derivatization

Solutions and reagents were brought to room temperature (RT) before initiating the extraction process. Standards (STD), sub-stocks, quality controls (QCs), surrogate matrix, and unknown human plasma samples are thawed at RT. Two double blanks are included in each batch by transferring 50μL of 2.5% BSA into clean 2mL microcentrifuge tubes. STD, QCs, and unknowns (50μl) were added to separate microcentrifuge tubes. Liquid chromatography mass spectrometry grade water (100μl) and 4% TCA (100μl) were added to these tubes and vortexed for 1 minute on a multi-tube vortex mixer (VWR International LLC, Radnor, PA).

Samples were centrifuged (Eppendorf 5415D) for 15 minutes at 10,000x g a 5°C. To a clean microcentrifuge tube, 200μL of LCMS grade water and 125μl 1M Sodium carbonate buffer were added. Each sample (125μL) was transferred to a microcentrifuge tube containing LCMS grade water and 1M sodium carbonate buffer. To these tubes, 25μL isobutyl chloroformate was added. All samples were vortexed for approximately 1 minute in a multi-tube vortex. The samples were then incubated for 15 minutes at 35°C and shaken at 150 rpm. Next, the samples were placed in the centrifuge for 10 minutes at 10,000x g at 5°C. After removing from the centrifuge, the samples were extracted using SPE cartridges (Waters Oasis HLB 10mg 1CC Cartridge) in conjunction with the UTC Positive Pressure manifold. Using low positive pressure, the columns were conditioned in the following sequence: Add 1mL of Methanol and wait until liquid flowed through, next add 1mL of LCMS grade water and wait until the liquid flowed through. Finally, samples (275μL) were loaded onto the cartridge. Samples sit for approximately 1 minute or until the sample flowed through entirely. Low positive pressure is applied to remove any residual liquid. Samples were eluted by adding 250μL 80:20:0.1 ACN:H2O:FA to each cartridge. Samples flow through with gravity, approximately 2 minutes, before using low positive pressure to elute the remaining liquid into clean microcentrifuge tubes. The eluant was dried under a gentle stream of N_2_ using a Turbovap at 37°C. All samples were reconstituted by adding 200μL of Reconstitution Solution (10:90:0.1, ACN:H2O:FA) and vortexed for 1 minute. Samples were then transferred to amber glass HPLC vials with 0.3 mL inserts. Samples were loaded directly for injection onto the LC-MS/MS, or the extracts were stored at 4°C until injection.

### 2.4 LC-MS/MS (MRM) Analysis for NAP

The multiple reaction monitoring (MRM) analyses was performed on an AB SCIEX QTRAP®5500 mass spectrometer (SCIEX) equipped with an electrospray source, Shimadzu (Kyoto, Japan) Ultra-Fast Liquid Chromatograph (UFLC) (LC-20AD XR pumps and SIL-20AC XR autosampler), and a Poroshell 120, EL-C18, (2.1 × 50mm, 2.7μ) column (Agilent, Santa Clara, CA). The MRM of derivatized N-Acetyl putrescine precursor and transition were m/z 231.00 and m/z 115.00, respectively, used as quantifier (Suppl. Table 1). Liquid chromatography was carried out at a flow rate of 0.350 mL/min, and the sample injection volume was 5 μL. The column was maintained at a temperature of 40°C. Mobile phase A consisted of 0.1% Formic Acid (Sigma Aldrich) in water (Fisher Scientific), and mobile phase B consisted of 0.1% formic acid in acetonitrile (Fisher Scientific). The gradient with respect to %B was as follows: 0 to 4 min, 5%; 4 to 5.5 min, 40%; 5.5 to 6.1 min, 95%; 6.1 to 10.0 min 5%. The instrument parameters for the 5500 QTRAP mass spectrometer were as follows: ion spray voltage of 5500V, curtain gas of 20 psi, collision gas set to “medium”, interface heater temperature of 500°C, nebulizer gas (GS1) of 40 psi, and ion source gas (GS2) of 40 psi and unit resolution for both Q1 and Q3 quadrupoles. Data analysis was performed using the AB SCIEX Analyst®software (version 1.5.1 or 1.6.2, Sciex, Framingham, MA), and peak integrations were reviewed manually.

### 2.5 Multi-Omic And Statistical Analysis

A multi-omic assessment of plasma was performed as previously reported [19]. The statistical analysis was conducted using R-Studio (2020, Version 3.6.2). Logistic regression was used to build all the models in ROC analysis. The selection of clinical variables was based on causal graphs (networks) generated by BERG’s AI platform bAIcis®, which relies on Bayesian network methods to learn from directed acyclic graphs [20]. To identify potential causal drivers of the PD status, an ensemble model from all variables was generated using bAIcis®, and the clinical variables directly connected to the outcome of interest were selected for further exploration. Variables of the best combination of multivariate models were chosen based on balancing the AUC and the complexity of the model. The 95% confidence interval was computed with 2000 stratified bootstrap replicates. Differences in means between PD and non-PD controls were assessed using t-test. Statistical significance for all analyses was determined at p<0.05.

## 3. Results

### 3.1 General Demographics and Discovery Assessment

A total of 201 PD patients and 199 non-disease controls participated in this study, with an average age of 65.4 and 64.7 years old, respectively (Table 1). The average score of Hoehn and Yahr (H&Y) scale was 2.1, including the 43 PD patients for stage I, 127 for stage II, 22 for stage III, 7 for stage IV, and 1 for stage V. The Unified Parkinson Disease rating scale (UPDRS) score for these PD patients ranged from 10 to 115. Metabolomic, lipidomic, and proteomic assessment of plasma samples were performed and integrated with Bayesian analysis and regression assessment of clinical and molecular features (Figure 1).

**Figure 1.**
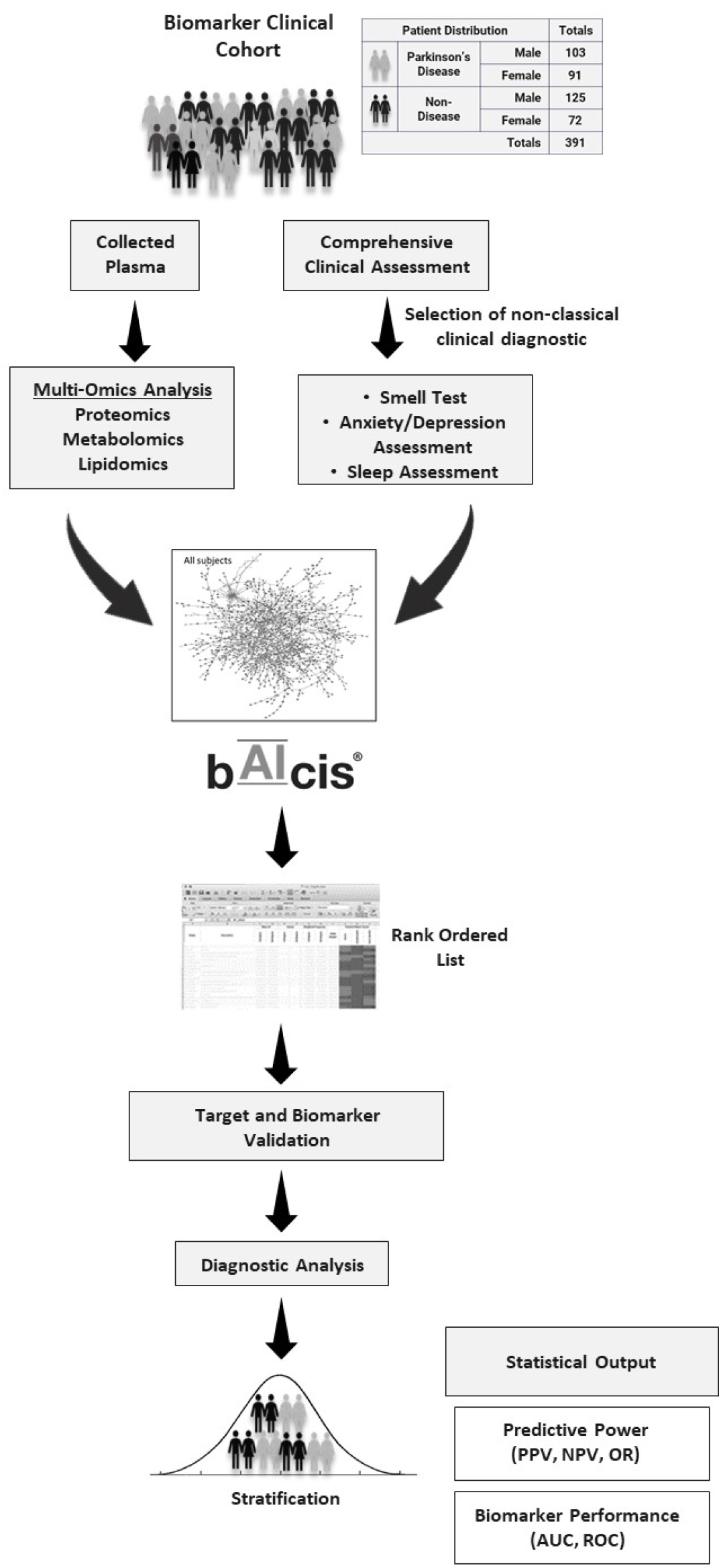
Study design. Biomarker Discovery Pipeline and Study Design. Single center observational study to assess markers in Parkinson’s patients and non-disease controls. Multi-omics analysis was performed and CLIA validated procedures were subsequently employed for quantitative biomarker assessment.

### 3.2 Assay Development and Validation

To quantify NAP in human K_2_-EDTA plasma, the quantification method using LC-MS/MS was developed. Due to low circulating levels of NAP, the sensitivity of quantification was improved using derivatization of isobutyl chloroformate in the sample preparation. The validation performance of the NAP assay is summarized in Suppl. Table 1–8, including linearity, precision, matrix effect, system suitability, short-term stability, long-term stability, and reproducibility in the autosampler. Fit-for-purpose method validation results demonstrated quantitative ranges for NAP from 1 ng/mL to 85 ng/mL in plasma analysis (Suppl. Table 3). The results of validation assessed by QCs met acceptance criteria (Suppl. Table 2–8).

### 3.3 Human Plasma Sample Analysis of NAP

Utilizing our CLIA validated quantification method, the K2-EDTA plasma samples from a total of 400 participants, including 199 non-disease (ND) and 201 PD cohort, were analyzed for NAP quantitation. When comparing PD in male and female patients, there were no statistical differences of plasma levels of NAP in gender that was observed (one-way ANOVA, p=0.6492) (Data not shown). A significant difference in plasma NAP between ND and PD was detected (t-test, p< 0.0001). The mean concentrations of NAP in the plasma for ND and in the PD cohorts were 3.70 ng/mL and 4.74 ng/mL, respectively (Figure 2).

**Figure 2.**
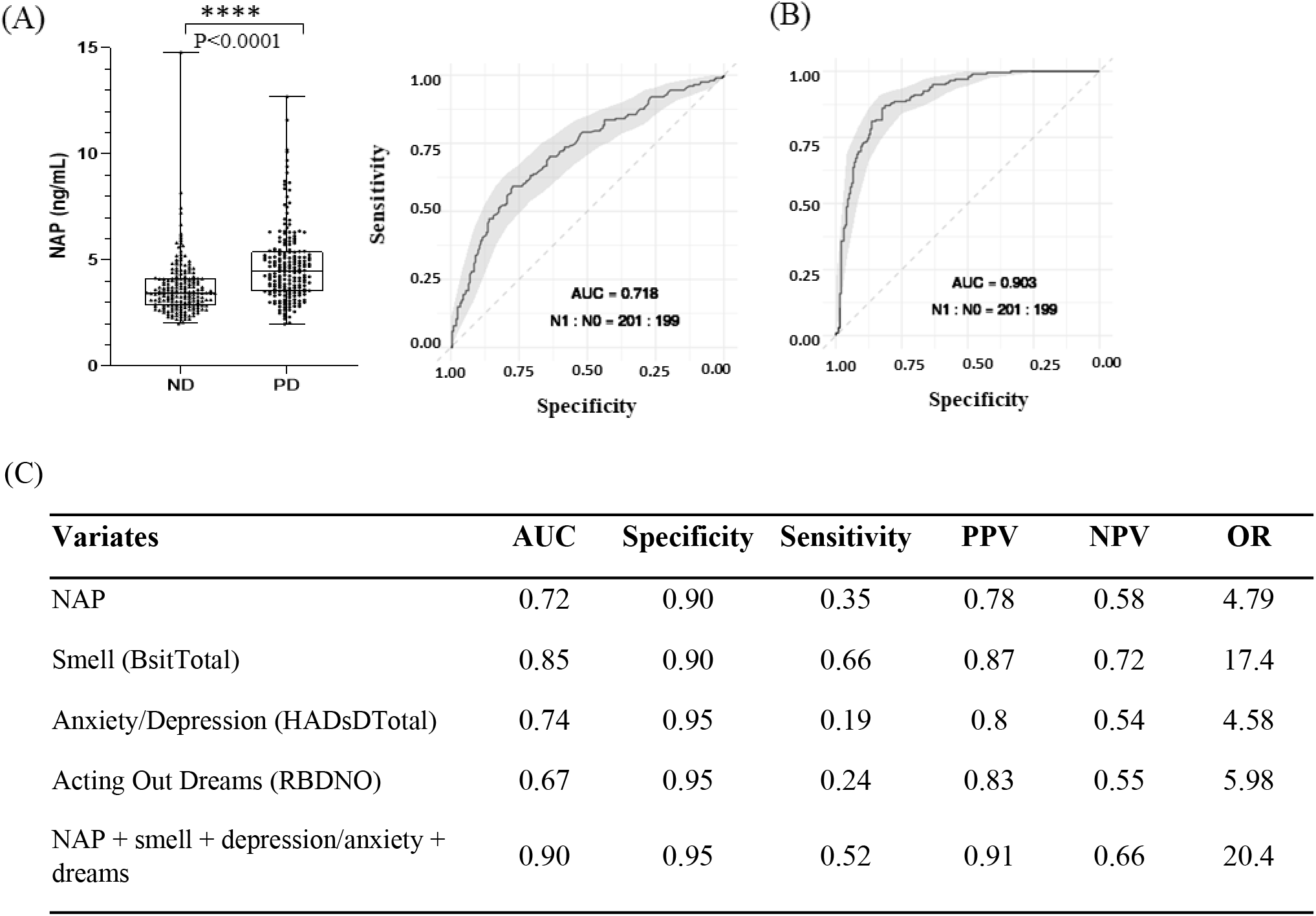
Plasma Sample Analysis for NAP and Receiver Operation Characteristic (ROC) Curve Analysis. (A) The plasma levels of NAP between non-disease and PD cohort.; ROC curve analysis for NAP alone (B) ROC curve analysis for NAP plus three clinical variables (C) Summary table for clinical performance of marker panel alone and combination, including area under curve (AUC), sensitivity, specificity, positive predictive value (PPV), negative predictive valve (NPV), and odds ratio (OR). 95% confident interval (CI) using Bootstrapping approach in ROC curve. Statistics was calculated by t-test, statistically significant: **** p<0.0001

### 3.4 Receiver Operating Characteristic Curve (ROC) Analysis of NAP and Three Clinical Features for PD Diagnosis

To examine the clinical performance, a receiver operating characteristic curve (ROC) analysis was applied for PD diagnosis in this project. In results of ROC analysis using the trapezoidal rule, the area under the curve (AUC) for NAP alone was 0.72, suggesting that plasma NAP levels demonstrate utility for PD diagnosis. The PD diagnosis of NAP alone showed the specificity as 90% and sensitivity as 35% with a cutoff value of 6.91ng/mL, respectively. In multivariate logistic regression models, the AUC values using the demographic factors of age, gender, or a combination with NAP were not improved in the separation of ND and PD cohorts (data not shown). Analysis of 121 clinical variables using the b<u>AI</u>cis platform identified smell test (B-SIT), depression and anxiety assessment test (HADS), and acting out dreams test (RBD1Q) as three potential clinical features for diagnosis of PD [16–18]. The multivariate model integrated with NAP and three clinical features revealed an optimal AUC of 0.90 to distinguish the PD from the non-disease cohort with a specificity of 95% and sensitivity of 52%. In predictive proficiency of PD diagnosis, the positive predictive value (PPV) and negative predictive value (NPV) for NAP alone were 0.78 and 0.58, and for the multivariate model, they were 0.91 and 0.66, respectively.

## 4. Discussion

Our study is the largest metabolomic biomarker study that identified a combination of clinical nonmotor measures (smell, anxiety/depression, and RBD) together with a plasma metabolite NAP with a specificity of 95% and sensitivity of 52%. It is also the first study that includes sufficient data for the calculation of diagnostic performance. PD is the second most common neurodegenerative disease affecting millions of people in the USA and many more worldwide. PD is estimated to occur in about 1% of the population over 60 and 4% of the individuals over 80 years old **[21, 22]**. It is difficult to accurately determine the precise prevalence of PD since the numbers do not include the majority of undiagnosed or misdiagnosed cases. The annual costs incurred for PD in the United States have been estimated to be nearly $11 billion, including $6.2 billion in direct costs **[23]**. The most significant proportion of cost for PD treatment occurs in the later stages of the disease when symptoms are most severe **[24]**. Any therapeutic strategy that could halt PD symptoms in the earlier disease stages with no further progression would greatly reduce disease burden for patients and families. In parallel, there is a critical need to develop diagnostic biomarkers for early disease detection using combinatorial PD biomarkers of clinical signs and blood metabolites.

Comprehensive understanding of human health and disease requires interpretation of complex biological processes at multiple levels such as genome, epigenome, transcriptome, proteome, and metabolome. These together can be classified as “multi-omics” data. The availability of multi-omics data has advanced the field of medicine and biology [25]. In this study, we have taken an integrative approach that has combined multi-omics data in order to highlight the interrelationships of the involved biomolecules and their clinical phenotype in disease. [26].

In three recent PD biomarker metabolomic analyses, several altered metabolites were identified, including amino acids, acylcarnitines, and polyamines in PD, however, these studies did not utilize CLIA-validated assays and were underpowered (Table 2). This first study used urinary metabolomic profiling of 18 metabolites, most of these branched chain, tryptophan, and phenylalanine amino acids, demonstrated discrimination capability in early, mid, and advanced stage PD patients (ND=65;PD=92) [27]. A major pitfall of the use of amino acids as biomarkers is that they could be overrepresented due to unbalanced demographics within the cohort or influence from concomitant medications. The second study assessed plasma metabolomic profiling and found acylcarnitines related to mitochondrial beta-oxidation as potential early diagnostic PD biomarkers. A group of long-chain acylcarnitines (AC12-14) (ND=32,45; PD=109,145) and nine fatty acid (8-18) metabolites (ND=40; PD=41) were found to be present in early stages of PD diagnosis [28, 29]. Due to the possible influence of other comorbidities within these groups, the use of acyl carnitines became a challenge for diagnosis. Moreover, increasing evidence suggests the possible role of polyamines in PD underlined several potential PD biomarkers in the mechanism of polyamine synthesis and metabolism [15, 30]. In a small scale clinical study (n=33), putrescine and N1-Acetylspermidine in cerebrospinal fluid (CSF) were significantly higher in PD patients compared to the control group [15]. Furthermore, the plasma level of N8-acetylspermidine and N1, N8-diacetylspermidine demonstrated significant accuracy in ROC curve analysis and positively correlated with H&Y stages [30]. Although these reports mentioned elevated NAP levels of plasma in PD patients, no further precise details of NAP clinical performance were elucidated. However, these measurements were not confirmed on robust bioanalytical platforms [30, 31].

**Table 2:**
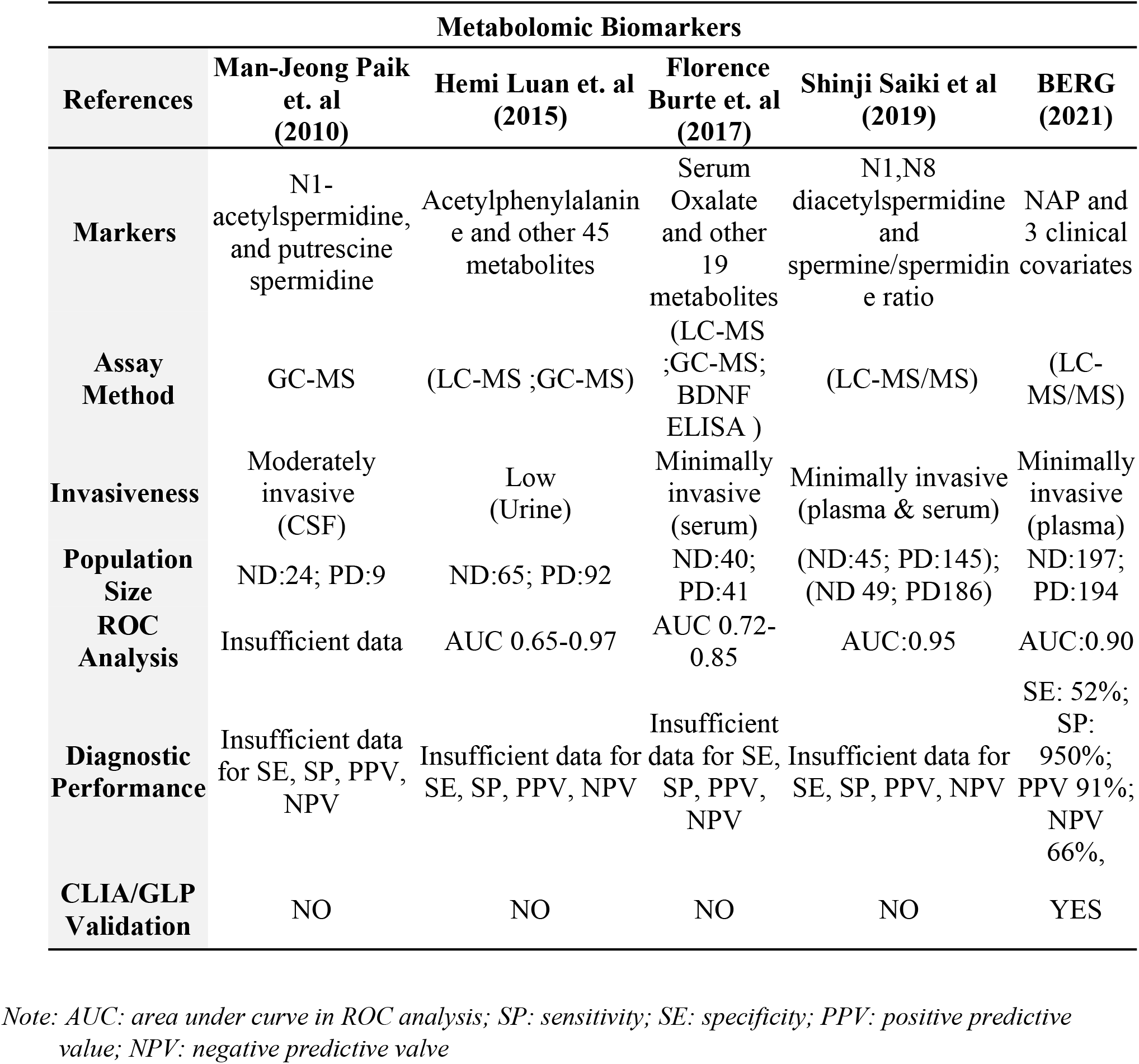
Summary of PD Biomarker Discovery from Published Literatures.

Some limitations in our study are listed as follows. While PD patients enrolled were not assessed by PET or SPECT imaging for the dopamine transporter (DAT) they were clinically diagnosed by a movement disorder specialist. Other illnesses and medication usage by patients may affect the status of their metabolism. Further validation of the proposed biomarker panel in more extensive, well-defined patient cohorts could be the next step to validate the finding of this study.

Lacking standardized criteria supporting PD diagnosis at the prodromal stage, current diagnosis of PD relies on primarily on clinical history and neurological assessment by a movement disorder and exclusion of other neurodegenerative diseases. The non-motor clinical signs and symptoms identified from our b<u>AI</u>cis® platform may appear in prodromal PD stage, including olfactory dysfunction, rapid eye movement sleep behavior disorder (RBD), and depression and anxiety. Three clinical features alone would be unreliable as a diagnostic biomarker [32, 33]. The multivariate logistic regression model with NAP and three clinical features demonstrated promising diagnostic performance. This study presents a potential avenue for clinical diagnostics while also underlining a possible role in polyamine metabolism for deciphering elusive etiology of PD.

## 5. Conclusions

In this study, we have completed the largest metabolite investigated biomarker study that was assessed using a CLIA validated assay and Bayesian analysis to identify complementary clinical features providing an AUC=0.9 and a PPV of 0.91. The multivariate marker model, consisting of three clinical features integrated with NAP, significantly improved the diagnostic accuracy for PD, demonstrating a higher positive predictive value than unintegrated individual variates. Clinical features alone demonstrated limited clinical efficacy. Results from this study demonstrated the power of a combined approach to PD biomarker discovery along with the possibility of implementing NAP into clinical biomarker tests.

## 7. Supplemental Materials

### 7.1 Assay Development and Validation of NAP

The following parameters were assessed during NAP assay validation:

#### 7.1.1 Calibration curve linearity

The linearity of eight independent calibration curves for NAP were assessed in human plasma assay.

#### 7.1.2 Intra and inter-batch precision

The assay was evaluated by analyzing the low limit of quantification QC (LLOQ-QC), low QC (LQC) medium QC (MQC) and high QC (HQC) human plasma (6 replicates for each) on different days.

#### 7.1.3 Matrix effect assay

Human K_2_-EDTA plasma and urine were spiked with the LLOQ-QC and HQC concentrations and assessed against calibration standards. Blank samples of plasma and urine were used as a zero standard from which a minimum of 9 different lots were extracted to determine the basal concentrations of NAP. These basal values were spiked by LLOQ-QC and HQC concentrations to obtain the actual nominal concentrations and then compared with the observed LLOQ-QC and HQC concentrations.

#### 7.1.4 Short-term stability (STS) and long-term stability (LTS) of calibrator in surrogate matrixes

Standard solutions for NAP were examined at the lowest concentrations (1.00 ng/mL) and the highest (85.0 ng/mL) in 2.5% BSA for the plasma assay, respectively. For STS, aliquots of calibrator were assessed by leaving both lowest and highest calibrators at 5°C for up to 24 hrs. These exposed samples were compared against unexposed samples. In LTS, aliquoted calibrators were stored at −80°C up to 88 days and evaluated by comparing these aliquots stored at −80°C against freshly prepared standards of the same concentration.

#### 7.1.5 STS, LTS, and freeze-thaw stability (FTS) of human plasma and urine samples

LQC and HQC samples were also used to assess for STS at RT for 24 hrs in the plasma assay, for LTS at −80°C up to 84 days, and for FTS up to four cycles at both −80°C and RT. In FTS, each freeze cycle was for a minimum of 24 hours. Triplicates or more for both QC samples were compared against the QC range generated during the validation.

#### 7.1.6 Re-injection reproducibility in the autosampler

Six replicates of each LLOQ QC, LQC, MQC and HQC samples were injected with a set of calibration standards. This same batch was re-injected, after being stored in autosampler at 4°C for up to 8 days for the plasma assay.

#### 7.1.7 Interference assessment

Potential interferences were evaluated by individually spiking LQC and HQC samples with human red cell hemolysate (500 mg/dL), unconjugated bilirubin (30 mg/dL), and triglycerides (1000 mg/dL) and then compared to the unspiked QCs in the plasma assay.

#### 7.1.8 System suitability, drift, and carryover

System suitability was assessed by calculating the precision (% CV) of the calculated concentration from 4 replicates at the beginning of each batch. The precision (% CV) was obtained by calculating concentrations from a minimum 4 replicates each of the system suitability standards at the beginning of the batch which must be ≤ 10%. Drift was assessed by calculating the %bias of the calculated mean concentrations for the system suitability standard from 4 replicates at the beginning of each batch and 2 replicate injections at the end of each batch. The % Drift between the SSS injections at the beginning and at the end of the batch must be ≤ 20%. In the carryover evaluation, a single injection of Reconstitution Solution after the System Suitability Standard was used to monitor carryover in each run. The carryover at the retention time and mass transition of NAP, must be ≤ 20% of the peak analyte area of lowest concentration standard.

**Supplemental Table 1:**
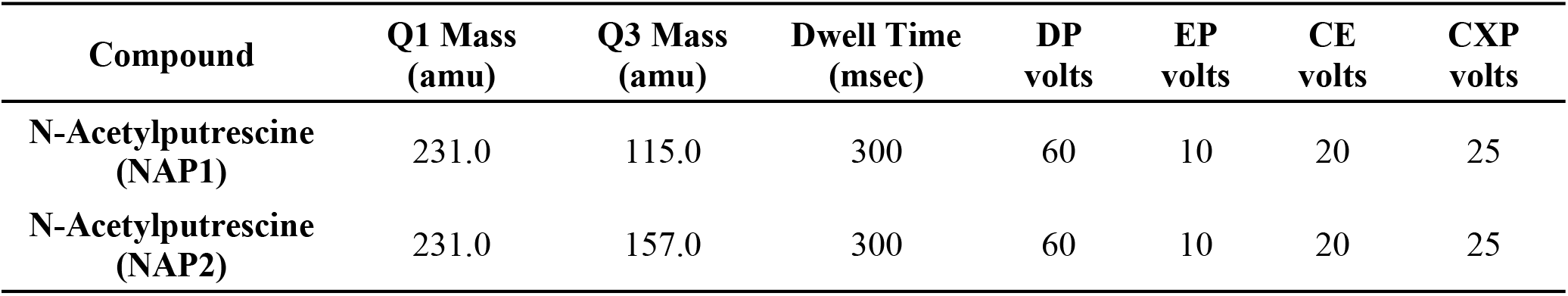
Summary Table of MRM Parameters

**Supplemental Table 2:**
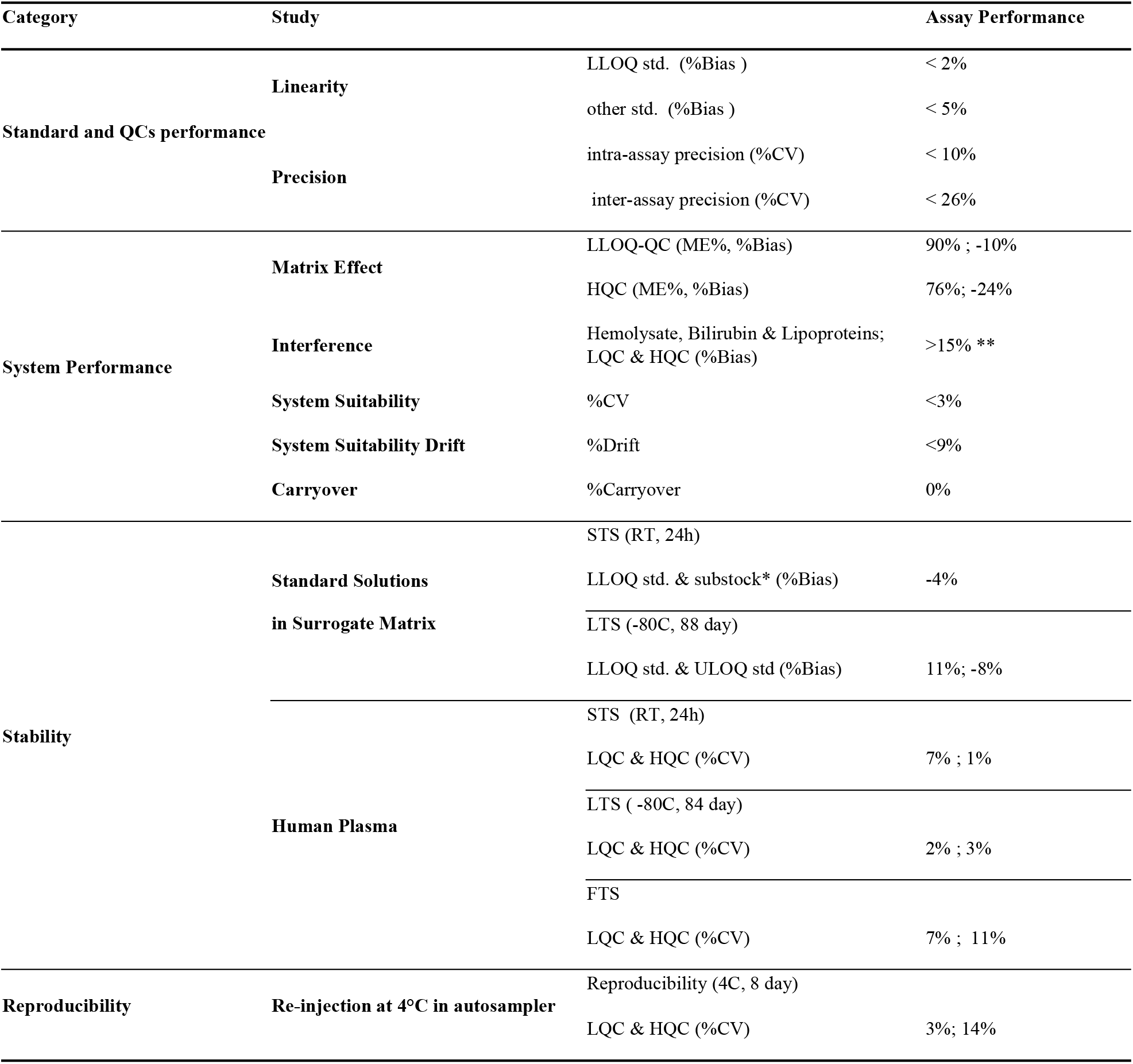
Validation Summary for NAP in Plasma Assay

**Supplemental Table 3:** Calibration Curve Ranges and Linearity for NAP in Plasma Assays.

| **NAP** |  | **Plasma assay (n =4)** |  |
| --- | --- | --- | --- |
| **Spiked Nominal**<br>**Concentration (ng/mL)** | **Mean Measured**<br>**Concentration (ng/mL)** | **%CV** | **Average %**<br>**Bias** |
| **1.0** | 1.0 | 3.5 | 1.5 |
| **2.0** | 1.8 | 5.2 | 4.1 |
| **5.0** | 5.0 | 6.6 | 2.4 |
| **10.0** | 10.1 | 3.1 | 2.0 |
| **25.0** | 23.9 | 3.9 | 2.0 |
| **50.0** | 52.4 | 2.7 | 2.6 |
| **68.0** | 70.8 | 6.0 | 1.4 |
| **85.0** | 83.4 | 4.5 | 2.8 |

Note: The linearity for NAP ( $r^2= 0.99745$ , $n=4$ )

**Supplemental Table 4.** The Intra- and Inter-Assay Precision (%CV) in Average for Six Batches

| **QC** | **Intra-assay**<br>**(% CV)** | **Inter-assay**<br>**(% CV)** |
| --- | --- | --- |
| **LLOQ QC** | 9.8 | 14.7 |
| **LQC** | 5.2 | 8.8 |
| **MQC** | 4.8 | 7.4 |
| **HQC** | 9.5 | 25.6 |

**Supplemental Table 5:** Matrix Effect Summary for NAP in Plasma Assay

| spiked with NAP in plasma (n=9) |  |  |  |
| --- | --- | --- | --- |
|  | spiking (ng/mL) | ME% | % Bias |
| LLOQ QC | 1.0 | 89.6 | -10.4 |
| HQC | 64 | 75.9 | -24.1 |

**Supplemental Table 6:**
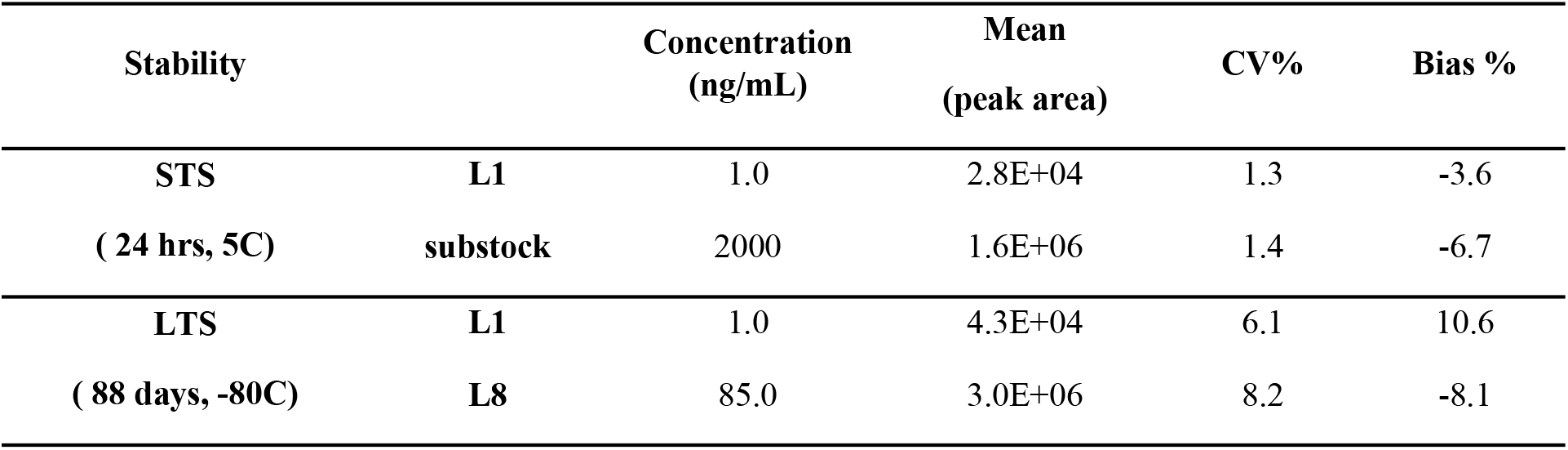
Short-Term Stability, and Long-Term Stability for NAP Spiked in 2.5% BSA Standard Solution

| Stability |  | Concentration<br>(ng/mL) | Mean<br>(peak area) | CV% | Bias % |
| --- | --- | --- | --- | --- | --- |
| STS<br>( 24 hrs, 5C) | L1 | 1.0 | 2.8E+04 | 1.3 | -3.6 |
|  | substock | 2000 | 1.6E+06 | 1.4 | -6.7 |
| LTS<br>( 88 days, -80C) | L1 | 1.0 | 4.3E+04 | 6.1 | 10.6 |
|  | L8 | 85.0 | 3.0E+06 | 8.2 | -8.1 |

**Supplemental Table 7:**
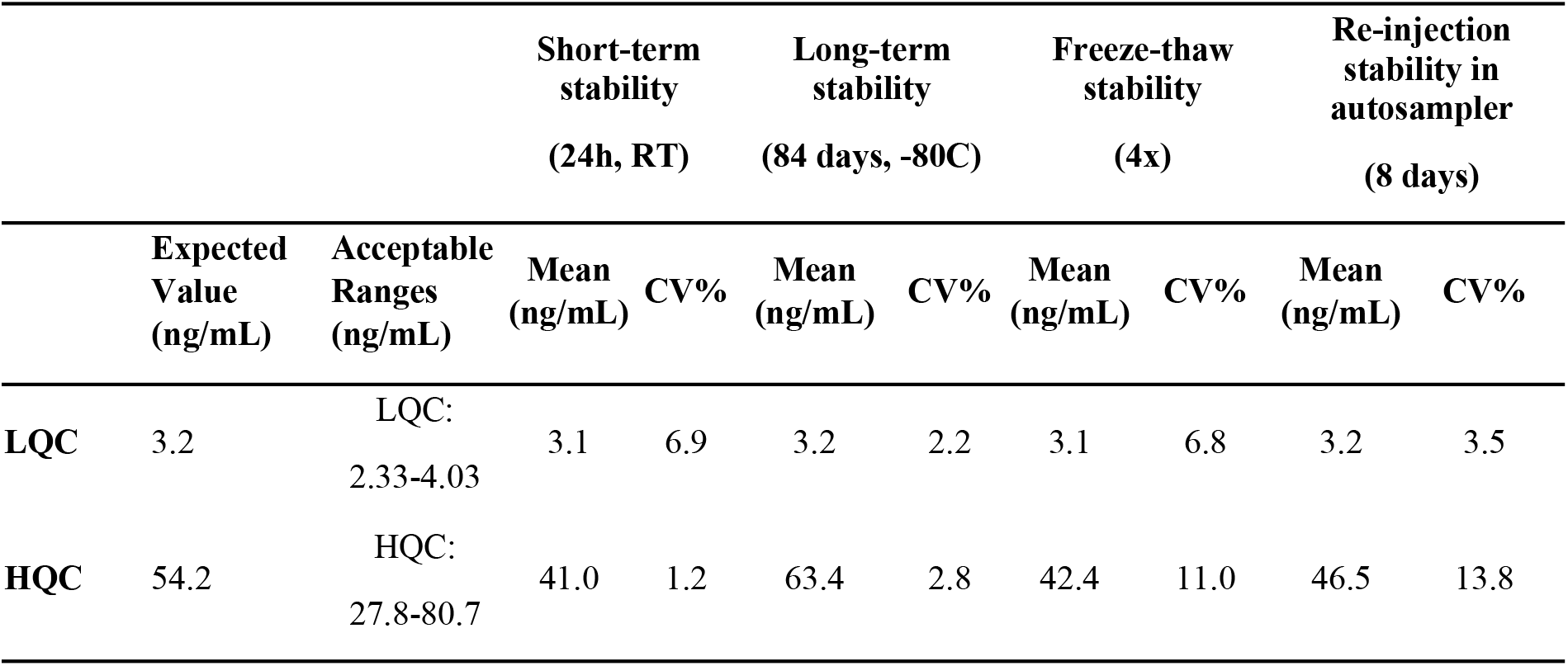
Short-Term Stability, and Long-Term Stability for NAP Spiked in Human Plasma

|  |  |  | Short-term stability<br>(24h, RT) |  | Long-term stability<br>(84 days, -80C) |  | Freeze-thaw stability<br>(4x) |  | Re-injection stability in autosampler<br>(8 days) |  |
| --- | --- | --- | --- | --- | --- | --- | --- | --- | --- | --- |
|  | Expected Value<br>(ng/mL) | Acceptable Ranges<br>(ng/mL) | Mean<br>(ng/mL) | CV% | Mean<br>(ng/mL) | CV% | Mean<br>(ng/mL) | CV% | Mean<br>(ng/mL) | CV% |
| **LQC** | 3.2 | LQC:<br>2.33-4.03 | 3.1 | 6.9 | 3.2 | 2.2 | 3.1 | 6.8 | 3.2 | 3.5 |
| **HQC** | 54.2 | HQC:<br>27.8-80.7 | 41.0 | 1.2 | 63.4 | 2.8 | 42.4 | 11.0 | 46.5 | 13.8 |

**Supplemental Table 8:**
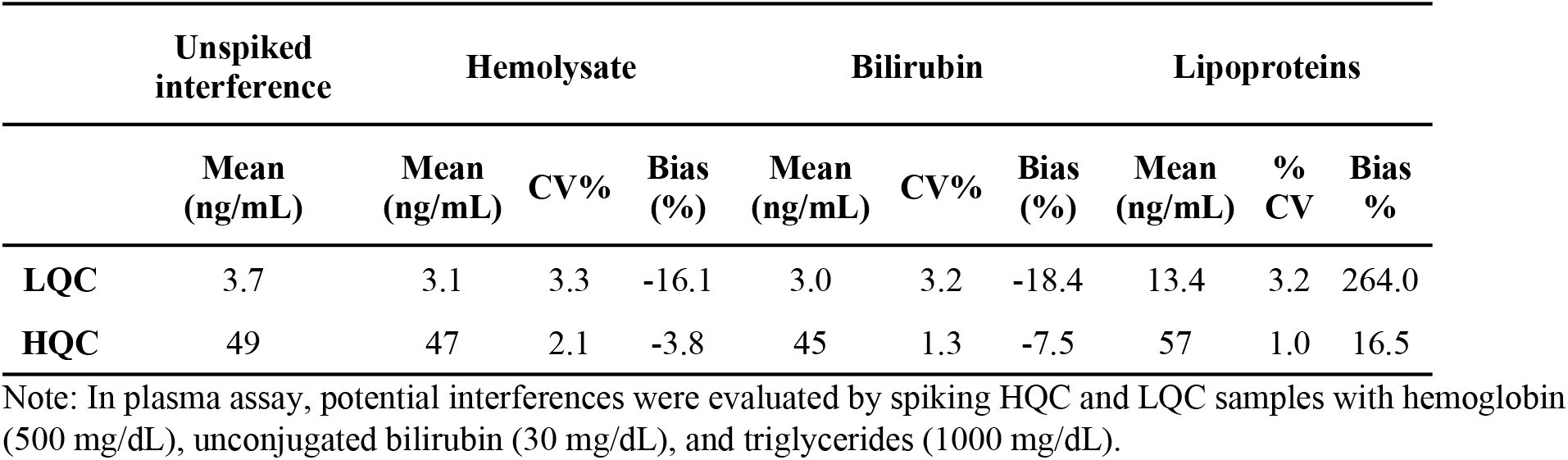
Interference in Plasma Assay

## Notes

### Competing Interest Statement

The authors have declared no competing interest.

